# Kmer-db: instant evolutionary distance estimation

**DOI:** 10.1101/263590

**Authors:** Sebastian Deorowicz, Adam Gudys, Maciej Dlugosz, Marek Kokot, Agnieszka Danek

**Author notes:** The authors contributed equally to the article (Sebastian Deorowicz, Adam Gudys).

## Abstract

**Summary:** Kmer-db is a new tool for estimating evolutionary relationship on the basis of k-mers extracted from genomes or sequencing reads. Thanks to an efficient data structure and parallel implementation, our software estimates distances between 40,715 pathogens in less than 4 minutes (on a modern workstation), 44 times faster than Mash, its main competitor.

**Availability and Implementation:** https://github.com/refresh-bio/kmer-db

**Supplementary information:** Supplementary data are available at publisher’s Web site

## 1 Introduction

Large volumes of data generated during the course of sequencing thousands of different organisms (100K Pathogen Genome Project (Weimer *el al.,* 2017), NCBI Pathogen Detection (https://www.ncbi.nlm.nih.gov/pathogens)), require fast analysis methods. Short substrings of nucleotide sequences, called *k*-mers, are commonly used in this area as they can be extracted either from genomes or sequencing reads, allowing assembly-free approach. They enable accurate approximation of evolutionary distances between organisms, thus are used for phylogeny reconstruction (Mash (Ondov *et al.,* 2016)), bacteria identification (StrainSeeker (Roosaare el *al.,* 2017)), or metagenomic classification (MetaCache (Müller el *al.,* 2017)). Importantly, if genomes are closely related, small subsets of *k*-mers are sufficient for obtaining acceptable accuracy, significantly reducing processing time. Nevertheless, asthenum-ber and the diversity of sequenced genomes continuously increases, the throughput of existing algorithms will soon become a bottleneck.

We introduce Kmer-db, a tool for *k*-mer-based analysis of large collections of sequenced samples. Thanks to a novel compressed *k*-mers representation and parallel implementation, our software is able to process thousands of bacteria genomes in minutes on a modern workstation.

## 2 Methods

As an input, Kmer-db takes *k*-mers extracted with KMC software (Kokot et *al.,* 2017) either from assembled genomes or read sets. The *k*-mers can be optionally filtered with a use of minhash (Broder, 1997) to save memory and time at the cost of accuracy.

The main analysis starts from *build* step, i.e., construction of a database for a set of samples on the basis of *k*-mers. A naive approach could be storing for each sample the corresponding fc-mer set. Excessive time and memory requirements make this representation prohibitve for large sample sets, unless fc-mer filtering method is used. Presented strategy is different. It is based on *k-mer templates,* i.e., lists of sample ids (s_id_). Such a list is defined for each fc-mer. The idea behind is that multiple *k*-mers may occur in exactly same samples, thus they share a template. Moreover, templates are often similar which allows further compression. As a result, Kmer-db consists of two basic structures: (i) a hashtable *K2T* mapping *k*-mers to corresponding template ids (t_id_), (ii) a table *CT* of compacted templates.

Samples are added to the database incrementally, with increasing identifiers. Let S indicate analyzed sample identified by s_id_. For each fc-mer from S, we record corresponding template identifier, t_id_, from *K2T* in an auxiliary array A (*k*-mers not present in *K2T* are inserted with special value t_id_ = 0). Array A is used to determine whether all *k*-mers with particular t_id_ are present in sample S. If so, template t_id_ from *CT* is extended with s_id_. If not, a new template is added to *CT* and corresponding entries in *K2T* are updated. The new template should contain all samples from considered t_id_ and additionally s_id_. To reduce redundancy, CT is a hierarchical structure—a new template stores only s_id_ on its list together with an identifier *p_id_* to its parental template. For this reason *CT* is referred to as *compacted templates.* Since samples are added to the database with increasing identifiers, lists of s_id_ in *CT* table are also increasing, thus they can be stored with a use of Elias gamma code (Elias, 1975), with about an order of magnitude space reduction compared to storing plain ids. The state of Kmer-db structures after adding five samples is presented in Figure 1. The intermediate states can be found in Supplementary Figures 1-5.

**Fig. 1.**
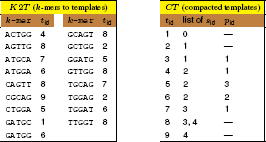
Database state after adding five samples: ACTGGATGCAG, GCTGGATGGAG, ACTGGATGGAG, ATGCAGTTGGT, CGCAGTTGGT. The structures can be used for obtaining list of samples for given fc-mer. E.g., fc-mer GGATG is assigned with template (£_id_) 5, whose parent (p_id_) is template 3, whose parent is template 1. Thus, GGATG is present in all samples (s_id_) from templates {5, 3, 1}, which are {2, 1, 0}.

The complete database can be further used for estimating evolutionary relationship between samples by determining numbers of common *k*-mers.

One of the available Kmer-db modes is *all2all* which determines matrix of common fc-mer counts for all samples in a database. When tens of thousands of samples are analyzed, matrix M of common *k*-mers counts requires gigabytes of memory. Therefore, maintaining cache locality when updating M elements is of crucial importance. For each template t_id_ from CT, the algorithm iterates over its s_id_ list and generates a collection of (s_id_, t_id_) pairs, stored in a cache-fitting buffer. Then, groups of pairs with same first element are identified. Note, that each group corresponds to a single M row: first element of a pair (s_id_) is a row number, while second element (t_id_) points to a template, whose entries indicate columns. The groups can be used to increment corresponding elements of M by template cardinality.

An alternative mode is *one2all* which produces vector V of common fc-mer counts between new sample S’ and all samples in a database. For all *k*-mers from S’ the algorithm selects corresponding templates, using *K2T* hashtable, and updates V accordingly.

The output of *all2all* and *one2all* stages are textual files with numbers of shared *k*-mers between pairs of samples and the total numbers of *k*-mers in each sample. They can be used to calculate various distance measures, e.g., Jaccard index, Mash distance. This is made by the *distance* mode.

## 3 Results

The experiments concerned calculation of distances between 40,715 genomes from NBCI Pathogen Detection on the basis of 20-mers. Samples were sorted w.r.t. species tax id (see Supplementary Material for other orderings). As the main competitor we selected Mash (Ondov *et al.,* 2016) since it implements essentially the same strategy as used in the NBCI Pathogen Detection project. Mash was configured to use 10,000 *k*-mers per sample (sketch size parameter), while Kmer-db was run in two configurations: (i) with minhashing at 2%_*o*_ threshold (to retain approximately same number of *k*-mers as Mash), (ii) on full fc-mer set. To evaluate software scalability, the subsets of 1k, 2k, 5k, 10k, and 20k samples were randomly selected from the full dataset. Table 1 presents the results of determining distance matrix on the basis of filtered *k*-mers (Mash *dist* step; Kmer-db *build* + *all2all* steps). Detailed results are presented in Supplementary Material.

**Table 1.**
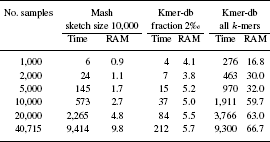
**Determining distance matrix on the basis of *k*-mers.**

Times are given in seconds, memory in GBs.

Kmer-db, when using 2%_*o*_ of *k*-mers was astonishingly fast. Evaluated on full dataset it was 44 times faster than Mash (212 s vs. 9,414 s) and needed less memory (5.7 GB vs. 9.8 GB). Analyzing all *k*-mers by Kmer-db (unfeasible to Mash due to computational requirements), took same time for all samples as running Mash on ~ 500 times smaller representation.

Importantly, our solution scaled well, especially in terms of memory usage. E.g., increasing sample set twofold from 20k to ~40k resulted in only 6% growth of RAM, which is thanks to the internal representation of Kmer-db (when database is large, a lot of *k*-mers from new samples share existing templates or their parts). We also noticed, that for increasing sets of samples, execution time of Kmer-db became dominated by matrix estimation, i.e., *all2all* step (see Supplementary Tables 1-2).

The approximate times of calculating similarity vector between a new fc-mer set and the already-build database (Kmer-db *one2all* step; Mash *dist* step) were: 1 s (Kmer-db 2‰), 1 min (Kmer-db all) and 5 s (Mash).

## 4 Conclusions

Superior running times and scalability of Kmer-db opens new opportunities in *k*-mer-based estimation of evolutionary distances. Our algorithm analyzed resampled fc-mer set of 40,715 bacterial genomes in less than 4 minutes-two orders of magnitude faster than Mash, confirming the readiness of Kmer-db for processing much larger datasets which are to appear in near feature. Presented approach was also able to compare distantly related genomes with few *k*-mers in common, where minhashing is inaccurate. Kmer-db was able to process all *k*-mers of analyzed bacteria in a time needed by the competitor for 500 times smaller fc-mer set.

## Funding

This work was supported by National Science Centre, Poland under projects DEC-2015/17/B/ST6/01890, DEC-2016/21/D/ST6/02952. The infrastructure was supported by POIG.02.03.01-24-099/13 grant: “GeCONil” Upper Silesian Center for Computational Science and Engineering”.

## Conflict of Interest

none declared.

